# Hybrid In Vivo Breast Cancer Model Reveals Transcriptomic Insights into Cancer Progression with Age

**DOI:** 10.1101/2025.08.27.672628

**Authors:** Lauren Hawthorne, Gokhan Bahcecioglu, Jun Yang, Emilija Aleksandrovic, Erin Howe, Siyuan Zhang, Pinar Zorlutuna

## Abstract

Aging is a key risk factor for breast cancer, yet the independent role of the extracellular matrix (ECM) in tumor progression remains understudied. Recent studies have investigated healthy mammary tissue and aging to understand their relationship; however, the independent effects of the aged ECM remain understudied. Herein, we describe a hybrid in vivo model where MCF10A ductal carcinoma (DCIS.com) cells – with or without knockdown of select targets – were seeded onto decellularized ECM from aged murine mammary glands and implanted into the mammary fat pads of young Rag1^-/-^ mice. Knockdown of selected targets, *IL1B* and *LOX*, reduced tumor growth on aged matrices in vivo and *P4HA1* knockdown enhanced tumor growth. Additionally, analysis of *LOX* on aged ECM highlighted *LOX* as a driver of tumor progression where knockdown reduced transcriptomic programs related to invasion and cellular stress. To isolate the individual ECM influence on tumor growth, MCF10A cells were seeded atop young or aged matrices where it was found that tumors grown on the aged ECM after implantation exhibited significantly greater volume and a larger tumorigenic region when compared to those from the young ECM. Furthermore, single cell RNA-sequencing revealed transcriptional enrichment of inflammatory and invasive genes within the aged matrix. Together, these results identify *LOX* as a driver of tumor progression and potential therapeutic target, and demonstrate that the aged ECM alone can promote breast cancer progression

## Introduction

Breast cancer incidence increases with age where it also disproportionately impacts older patients who experience worse outcomes.^1–4^ Despite advances in screening and treatment, it remains a prevalent and metastatic disease with an ever-increasing incidence rate, primarily attributed to the growing aging population and the associated biological mechanisms that drive changes in aged breast tissue properties.^1,2,5,6^

Historically, aging research has focused on genetic and cellular-level changes – such as genomic instability and senescence – many of which overlap with the hallmarks of cancer.^7^ This perspective has provided key insights into how aging influences cancer initiation and progression, leading to breakthroughs in areas such as oncogene activation, tumor suppressor loss, and DNA repair deficiencies in aged tissues.^7,8^ Other major changes observed linking aging and cancer haven been the extracellular matrix (ECM) and its composition and mechanics along with immune system dysregulation. The ECM is a dynamic structure that provides mechanical and biochemical cues to cells. As the body ages, ECM naturally stiffens due largely to the crosslinking of collagen. This stiffening of the matrix influences how cancer cells proliferate, invade, and respond to therapy. Additionally, both aging and cancer lead to compositional changes that create an imbalance in the normal environment, thus promoting invasion and metastasis.^9 10–14^ In addition to mechanical and compositional changes, the immune system also becomes dysregulated within both the aged and cancerous environment through a process of immunosenescence.^15,16^ In this, some immune cells, such as macrophages, can become upregulated and tumor-associated where they begin to promote cancer-permissive factors that promote cancer cell invasion.^17–20^ Several recent studies linking aging and cancer have shown that the cellular population (epithelial, stromal, and immune cells) shows a shift in proportion and cellular identity, influencing cancer risk and the aged microenvironment.^21,22^

While aging is a well-established risk factor and previous studies serve as comprehensive source, the independent effects of the aged extracellular matrix (ECM) on cancer predisposition, onset, and progression remain understudied.^23,24^ While recent studies have looked into cellular population shifts due to aging, they overlook how age-related changes in the ECM directly influence cancer development, and those that do are unable to decouple other systemic aging effects, such as immunosenescence. Addressing this gap requires models that can isolate the ECM’s independent contributions to tumor growth. While most breast cancer age-related models rely on comparisons between young and aged animals, the length of time required to develop an appropriately aged animal that mimics human age is substantial – increasing the financial and material costs.^25–27^ Additionally, these studies typically do not decouple the age-related alterations in the ECM from other systemic differences, including cellular senescence and immunosenescence, also known to increase susceptibility to cancer.^28^ These lead to paradoxical results where studies performed on aged mice have shown varied results, with some showing that with age, the ECM changes can lead to more invasive and prone to distant metastasis^29^, while others show that older patients have a slower tumor growth and reduced metastasis.^30^ Understanding these mechanisms, as well as the differences between in vitro and in vivo environments, is critical for identifying new therapeutic targets and evaluating the influence and effectiveness of existing and future treatments.

To address current limitations and better understand the mechanisms of breast cancer progression and metastasis and to enhance recent studies^21,22^, **we developed a hybridized model that allows for the independent study of the aged ECM and its role in breast cancer development and progression**. We first generated stable knockdown MCF10 DCIS.com cell lines of genes known to be enriched within both the aged matrix and breast cancer – *LOX, IL1B*, and *P4HA*. In vivo studies utilizing showed that *LOX* and *IL1B* contributed to a decrease in tumor growth, while *P4HA1* knockdown resulted in an increase in tumor volume. When comparing the *LOX* knockdown group to the control group in vivo, scRNA sequencing results showed that knockdown-suppressed genes are responsible for mostly tumor progression and metastasis. Moreover, *LOX* knockdown inhibited gene clusters responsible for tumor-promoting immune response, suggesting roles in the immune response. After validating the differential response within our model, we moved to assess the effect that ECM alone influences upon cancer growth and development. We find that the aged ECM alone was sufficient to enhance tumor growth within the same-host model. Transcriptomic analysis through single-cell-sequencing further showed that the aged ECM upregulates genes related to cancer cell initiation, progression, and metastasis while downregulating genes related to ECM stability, cell signaling, and cancer suppressive immune response.

## Results

### Knockdown of LOX and IL1B Suppress Tumor Growth on Aged ECM

To first investigate ECM-mediated tumor progression with age, we identified several targets known to literature to be upregulated within the aged environment and related to tumor growth and metastasis: *IL1B*^31,32^, *P4HA1*^33,34^, and *LO*X^13,35^. We first generated and validated stable knockdown cell lines (shIL1B, shLOX, and shP4HA1) from MCF10A DCIS.com cells (**Fig. 1A, 1C, 1E**). From this, we engineered a novel hybridized mouse model by utilizing decellularized ECM (dECM) derived from aged (24-48 months) murine mammary glands of C57BL/6 mice and implanted them into young Rag1^-/-^ mice. Before implantation, dECM were seeded with a control (shNT) and one of the knockdown cell lines. The mice were split into two groups with reversed implantation locations to eliminate the influence of environmental variation on tumor growth between the left and right sides of the mammary fat pad (**Fig. 1A**). This experimental design allowed for a direct comparison of the tumor growth between control and knockdown tumors under controlled conditions. Each line was seeded onto aged dECM and implanted into young Rag1^-/-^ mice mammary fat pad, with matched shNT controls implanted on the opposing side. After 6 weeks, mice were sacrificed, and tumors were excised for analysis. shLOX tumors were significantly smaller than shNT (n = 9, p = 0.0135). *IL1B* knockdown resulted in a non-significant decrease in tumor volume compared to control (**Fig. 1D**). Interestingly, *P4HA1* knockdown increased tumor volume compared to control, suggesting a potentially context-dependent role in the aged ECM (**Fig. 1F**)

**Figure 1:**
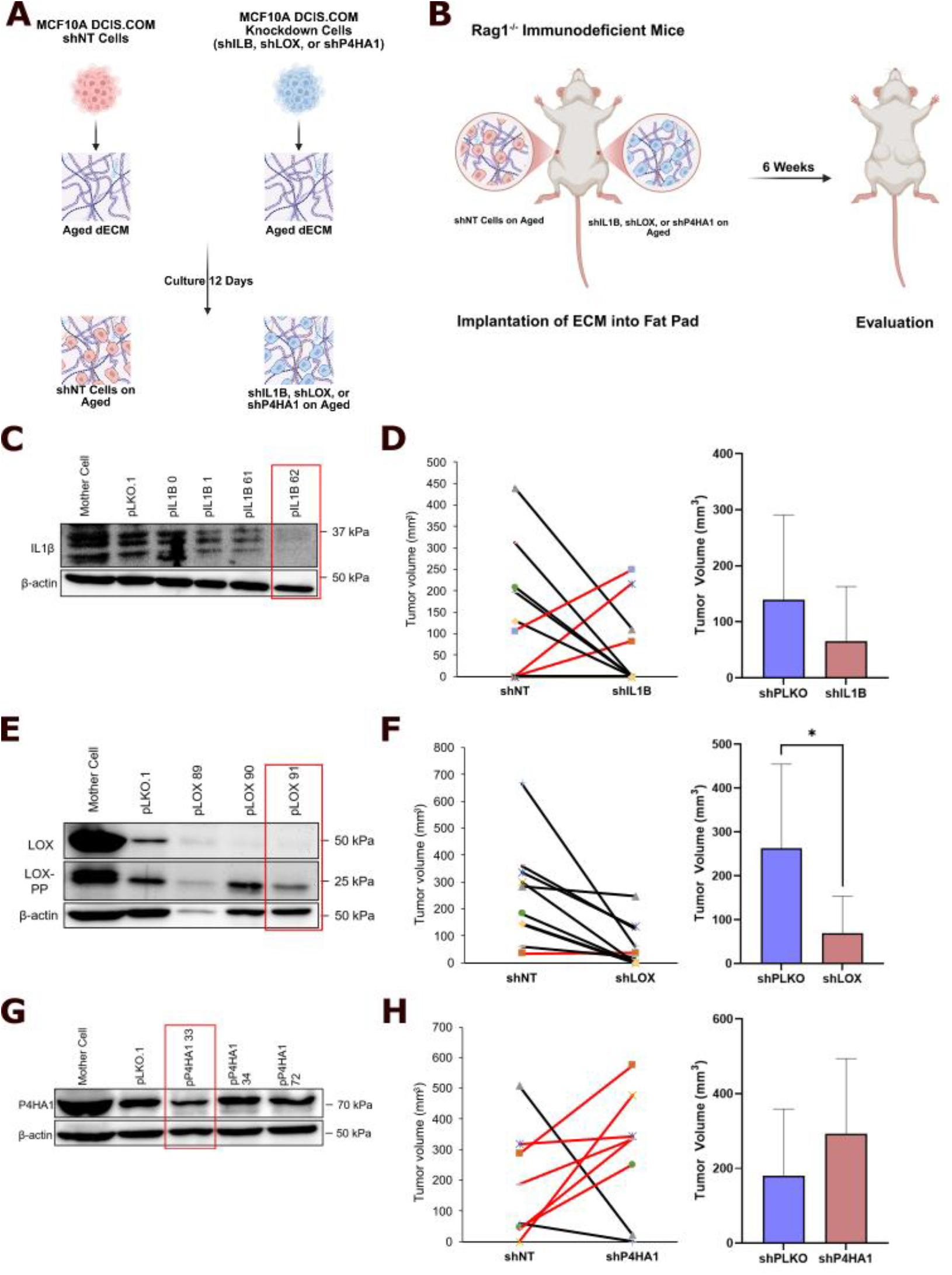
*LOX* knockdown suppresses tumor growth in aged ECM. **(A)** Representative schematic of shNT or knockdown cell seeding atop aged dECM. **(B)** Representative schematic of implantation of recellularized aged ECM into immunodeficient mice. **(C)** *IL1B* knockdown confirmed by Western Blot. **(D)** Tumor volumes of shNT control vs. shIL1B group (p > 0.05, n = 10). **(E)** *LOX* knockdown confirmed by Western Blot. **(F)** Tumor volumes of shNT control vs. shLOX group (p = 0.0135, n = 9). **(G)** *P4HA1* knockdown confirmed by Western Blot. **(H)** Tumor volumes of shNT control vs. shP4HA1 group (p > 0.05, n = 8).

### LOX Knockdown Suppresses Pro-Metastatic Cell State and Gene Expression

As *LOX* was found to be the most impactful and consistent on tumor development and growth, we next processed shLOX and their respective shNT tumors for scRNA-sequencing (scRNA-seq) analysis to further study the differences at a transcriptomic level. Before scRNA-seq processing, CD11b^+^ (myeloid cells) and CD11b^-^ (tumor and stromal cells) were separated using magnetic beads enrichment and sequenced separately. scRNA-sequencing was performed on both the tumor (**Fig. 2, Table S1**) and immune (**Fig. S1, Table S2**) cells.

**Figure 2:**
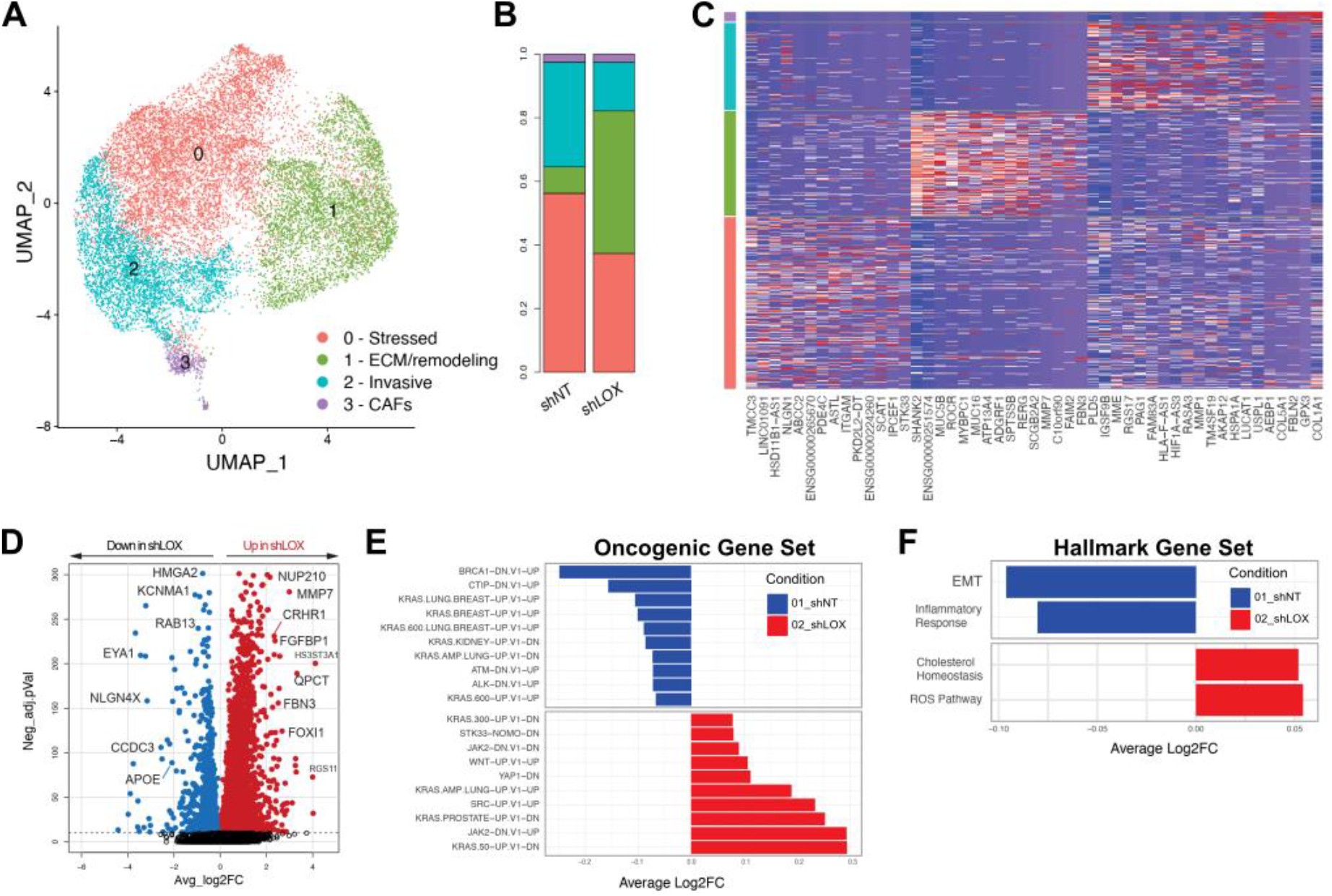
LOX knockdown suppresses invasive tumor cell states and oncogenic transcriptional programs. **(A)** UMAP of all cells captured, split by treatment and **(B)** bar chart depicting the ratio between the clusters in shNT and shLOX groups. **(C)** Heat map of top cluster marker genes for each identified cluster. **(D)** Differentially Expressed Gene (DEG)-derived volcano plot of gene level differences between shNT and shLOX and bar chart graphs for the GSEA-derived **(E)** C6 Oncogenic Signature Gene Set and **(F)** Hallmark Signature Gene Set

Analysis of the tumor cells revealed 4 distinct clusters and their respective marker and significant genes **(Fig. 2A-C)** – Cluster 0: stressed, Cluster 1: ECM/remodeling, Cluster 2: invasive, Cluster 3: cancer-associated fibroblasts (CAFs). LOX knockdown led to reduced relative proportions of Clusters 0 and 2 and an increase in Cluster 1. Cluster 0 exhibits high expression of *LINC01091* and *HSD11B1-AS1*, both of which are long noncoding RNAs, the latter having known regulatory roles in cancer.^36–40^ In addition, high expression of *ABCC2* and *TMCC3* have been linked to chemoresistance and stress response.^41–43^ Cluster 1 expresses markers such as *MUC5B, MUC16*, and *MMP7*, indicating that this cluster has ECM remodeling and mucin production roles.^44–47^ Cluster 2 expresses several invasion and migration related genes, such as RASA3 and MMP3,^48–50^ indicating a pro-metastatic program. In addition, this cluster is marked by immune evasion and hypoxic response genes such as *HLA-F-AS1* and *HIF1A-AS3*, respectively.^51–55^ Cluster 3 CAFs are marked by ECM genes such as *COL5A1, AEBP1*, and *FBLN2*. Within the shLOX group, the relative frequencies of Clusters 0 and 2 decreased, whereas the relative frequency of Cluster 1 increased (**Fig. 2B**). This shift suggests that after intracellular *LOX* knockdown, the transcriptional programs with roles in tumor progression and metastasis are suppressed, whereas those responsible for ECM structure are enhanced.

### LOX Knockdown Alters Oncogenic and Hallmark Signaling Pathways

The transcriptional differences between shLOX and shNT were further examined through differential gene expression (DEG) analysis (**Table S3**). Genes such as *HMGA2*^56^ and *KCNMA1*^57^, which have been shown to be linked to various cancer types as biomarkers for cancers initiation and invasion, were significantly downregulated on shLOX tumors (**Fig. 2D**). Gene Set Enrichment Analysis (GSEA) of the Oncogenic Gene Set (**Table S4-S5**) showed downregulation of *JAK, Wnt*, and *YAP* signaling – mechanosensitive pathways which are often dysregulated in cancerous environments.^58–60^ *BRCA1*^61^ and *ATM*^62^ – DNA repair pathways – were also downregulated. Interestingly, *KRAS*-related gene-sets were elevated in both shNT and shLOX groups, suggesting that *LOX* may play a dual role in modulating this pathway in the aged microenvironment (**Fig. 2E**). Further analysis with the Hallmark Gene Set (**Table S6-S7**) demonstrated suppression of epithelial-to-mesenchymal-transition (EMT) and inflammatory pathways while showing upregulation of pathways related to cholesterol homeostasis and reactive oxygen species (ROS) (**Fig. 2F**). Together, these results indicate that the knockdown of *LOX* is capable, within the aged environment, of suppressing pathways beneficial for tumor proliferation, invasion and stress response, supporting its role as a key effector of matrix-induced tumor progression.

### Aged ECM Drives Tumor Growth In Vivo

With observed differential tumor growth between control and knockdown cells lines in our model, we next aimed to isolate the independent contribution of the aged ECM to tumor progression. Rather than assessing with differential cell lines, we utilized decellularized ECM (dECM) derived from young (2-4 months) and aged (24-48 months) murine mammary glands of C57BL/6 mice and implanting them into young Rag1^-/-^ mice. Before implantation, dECM were seeded with normal MCF10A DCIS.com cells. The mice were split into two groups with reversed implantation locations to eliminate the influence of environmental variation on tumor growth between the left and right sides of the mammary fat pad (**Fig. 3A-B**). This experimental design allowed for a direct comparison of the tumor growth in young vs. aged environments under controlled conditions. After, tumors on the aged dECM were found to be significantly larger, with an average volume of 327 ± 132 mm^3^, compared to those on the young dECM, which averaged 162 ± 60 mm^3^ (p= 0.0032, n = 10) (**Fig. 3C-D**). Further histological analysis with hematoxylin and eosin (H&E) staining confirmed an approximate 14% average increase in tumorigenic region in tumors from the aged sides (**Fig. 3E-F**), confirming that the aged ECM independently promotes tumor growth and tissue invasion.

**Figure 3:**
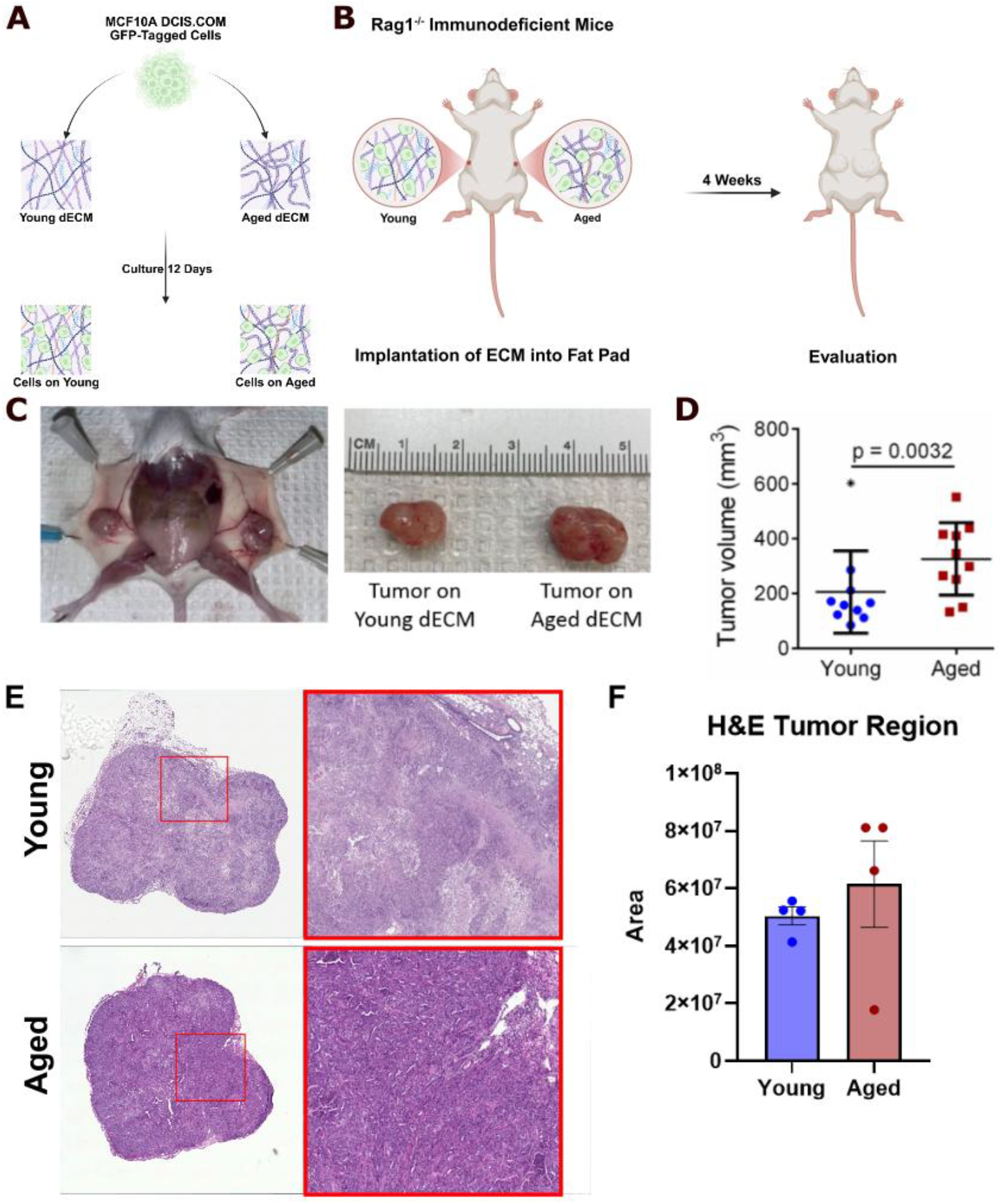
Aged ECM Independently Promotes Tumor Growth In Vivo: **(A)** Schematic of MCF10A DCIS.COM cells seeded atop aged or young dECM. **(B)** Schematic of implantation sites and subsequent tumor growth. **(C)** Representative images of tumors grown. **(D)** Quantification differences in tumor growth where aged ECM had a significantly higher tumor growth (p = 0.0032, n = 10). **(E)** Representative H&E images of tumors from aged or young ECM. (F) H&E quantification (n = 4, p > 0.05).

### Aged ECM Enriches Invasive and Inflammatory Phenotype

To better understand the underlying transcriptomic mechanisms and cell populations contributing to tumor growth and development between the groups, scRNA-seq was performed on the excised tumors (**Fig. 4, Fig. S2, and Table S8**). Transcriptomic analysis revealed 12 distinct cell clusters and their defining marker genes (**Fig. 4A-C**). These clusters exhibited a differential proportional distribution between tumors grown in young vs. aged ECM conditions.

**Figure 4:**
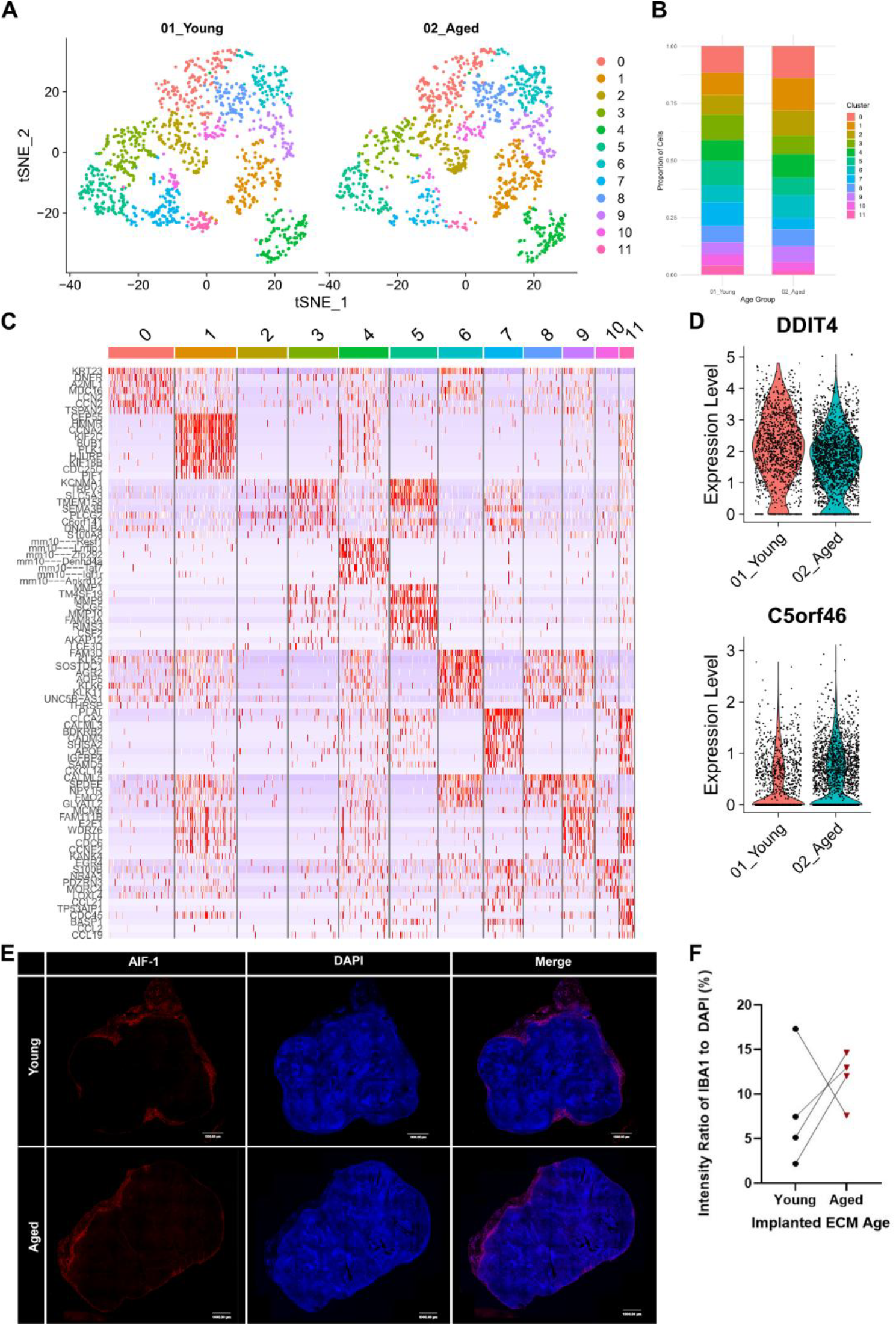
Aged ECM drives distinct transcription cell states in tumors. **(A)** tSNE of all cells captured, split by age and **(B)** bar chart depicting the ratio between the clusters in aged and young groups. **(C)** Heatmap of scRNA-seq gene expression reveals 12 tumor cell clusters with distinct transcriptomic profiles. **(D)** Distinct genes upregulated on the young and aged ECM. **(E)** Representative immunofluorescence of AIF-1 expression on young vs. aged tumors and **(F)** quantification.

Clusters 0, 1, and 11 were found to be most expanded within the aged matrices and were mostly related to ECM remodeling, cell cycle regulators, and genome regulators, respectively. Cluster 0 showed upregulation of genes such as *KRT23*^63,64^, *DNER*^65,66^, and *TSPAN2*^67,68^ which are typically associated with cell structure and various cellular processes. These genes have been previously implicated as cancer biomarkers where they play a role in cell migration, invasion, downstream gene expression, and metastasis. Cluster 1 displayed strong expression of mitotic and cell cycle regulators including *CEP55*^69,70^, *HMMR*^71,72^, and *CCNA2*^73,74^, as well as microtubule-associated proteins – such as *KIF2C*^75,76^ and *KIF18B*^77,78^. Together, these indicate a highly proliferative transcriptional state within the aged ECM. Finally, Cluster 11 was enriched in genes relating to DNA replication and cell cycle regulators such as *MCM6*^79,80^, *E2F1*^81,82^, *WDR76*^83,84^, and *DTL*^85,86^. This further suggests that the aged ECM promotes aberrant replication and proliferative signaling, with many of the upregulated genes having been previously linked to cancer-associated cell cycle dysregulation and proliferative biomarkers. Conversely, within the young ECM Clusters 3, 5, and 7 were found to be enriched with roles in ion channel and immune regulation, ECM stability, and tumor suppression genes. Cluster 3 was highly defined by the expression of various ion channel regulators such as *KCNMA1*^87,88^ and *TRPV3*^89,90^, alongside immune regulatory genes including *PLCG2*^91,92^ and *C6orf141*^93,94^, suggesting a role in maintenance of coordinated cellular communication and response and immune mediated tumor suppression. Cluster 5 exhibited high level of ECM stability and remodeling genes – *MMP1*^95^, *MMP9*^96^, *MMP10*^97^, indicating ECM remodeling within the young ECM. While these MMPs are frequently associated with cancer initiation and invasion^98–101^, in the young ECM they marked a state of remodeling redirected toward tumor suppression rather than initiation. Finally, Cluster 7 was enriched in tumor suppressor genes including *CLCA2*^102,103^, *CADM3*^104,105^, further supporting that the young ECM fosters a microenvironment that limits tumor progression. Complementing the cluster-specific findings, volcano plot analyses of significant genes (**Fig. S3**) revealed that the aged ECM biases dual-function genes toward tumor-suppressive programs. Notably, *DDIT4*^106^, typically characterized as a biomarker of cancer progression^107^ but also documented as a tumor suppressor^108^, was upregulated, demonstrating influence of the aged microenvironment’s role in tumor development. Similarly, the aged ECM showed enrichment of *C5orf46*^109^, a gene correlated with poor prognosis due to its role in driving malignant phenotypes and reshaping the immune landscape toward tumor support^110^ (**Fig. 4D**). To further contextualize these changes, we investigated *AIF-1* – a gene previously implicated in breast cancer development, progression, and metastasis – which revealed an average 12% expression in tumors gown on aged ECM, compared to 8% in young, within the same host model (**Fig. 4E-F**). Up-regulation of *AIF-1* suggests tumor cells adopt a unique inflammatory signature in the aged microenvironment. Together, the age-respective profiles suggest that the aged ECM promotes a tumor permissive environment, contrasting the young ECM’s immune regulation and tumor suppression.

## Discussion

This work is particularly critical as the aging ECM and its effects are becoming increasingly acknowledged as a key component in tumor progression. While several recent, high-resolution, single-cell sequencing studies have looked into mammary murine tissues revealing compositional shifts in epithelial, stromal, and immune cell populations in mice in both healthy and the tumor microenvironment with respect to aging^21,22^, these studies fail to decouple the independent effect of the ECM as a contributor to tumor growth and development.

Using knockdown and control cell lines implanted within the same host on aged ECM we first find that *LOX*, a key protein upregulated within the aged environment, emerged as a key regulator of ECM induced tumor progression. *LOX* knockdown significantly and consistently reduced tumor volume when compared to non-knockdown control cells within the same host. Further scRNA-seq analysis of the shLOX group on aged matrix revealed decreases in transcriptional programs related to cellular stress and invasion. We see a clear transcriptional shift away from pro-inflammatory and invasive programs and toward ECM structure and organization. Cluster 1, enriched after LOX knockdown, was characterized by genes such as *MUC5B, MMP7*, and *MUC16* – associated with mucin production and matrix regulation – indicating a more structurally engaged and less migratory. Complimentary to these transcriptional alterations, GSEA analysis at the pathway level revealed a down regulation of various mechanotransducive and oncogenic programs. The Hallmark Gene set revealed *LOX* knockdown significantly reduced pathways related to EMT and inflammation while increasing those related to cholesterol homeostasis and ROS. Additionally, analysis of the Oncogenic Gene Set showed *LOX* knockdown and its dual role in various pathways such as *KRAS*, upregulation of pathways such as *JAK* and downregulation of pathways such as *BRCA1*. These results highlight *LOX* as a potential therapeutic target that inhibits tumor growth by enhancing ECM dynamics and reducing metastatic potential. Additionally, it shows that *LOX* alone has the potential to facilitate tumor proliferation and development and that its cellular knockdown within the aged environment alone can significantly reduce the tumor volume.

Furthermore, we demonstrate that the aged ECM alone creates a tumor-permissive microenvironment that promotes tumor growth and metastasis. We find that the tumor growth within the aged ECM is significantly greater than that of the young, within the same host. H&E analysis further showed a greater cancerous region within the excised aged tumors when compared to their young counterpart. Transcriptional profiling confirmed that the aged ECM independently promotes an environment conducive to tumor growth and metastasis by enriching clusters associated with cell cycle activation, DNA replication, and chromatin regulation. Moreover, the aged ECM demonstrated enrichment of *C5orf46*, a gene associated with adverse outcomes by facilitating malignant progression and immune evasion, highlighting the tumor-permissive nature of the aged ECM. Similarly, the aged ECM, with its larger tumor volume, upregulated known tumor-suppressor and promoter gene *DDIT4*. The increased tumor size reflects context-dependent reprogramming driven by the aged ECM, which enhances the tumor promoting activity and demonstrates its capacity to independently influence cancer progression.

In contrast, the young ECM alone is indicated to promote a tumor suppressive microenvironment through the upregulation of ECM stabilizing and cell signaling genes. Tumors gown on the young ECM were transcriptionally enriched in ion channel regulators and immune-modulatory genes, which were indicative of a more stable and regulated microenvironment that supports cellular communication and suppresses tumor initiation. These findings indicate that the ECM age is a deterministic factor in shaping a tumor permissive environment, where the aged ECM can accelerate initiation and metastasis.

The aged ECM tumors exhibited an increase in unique inflammatory signals, seen by upregulation of *AIF-1*, an inflammatory regulator that is also implicated in breast cancer and linked to EMT and metastasis. This contrasts with the immune-regulator signals found within the young ECM tumors where the immune enrichment was marked by known, tumor suppressive genes *PLCG2* and *C6orf141*. Together, this work supports recent studies whereby the aging environment enhances tumor initiation and metastasis due to cell cycle and immune changes.^21^

These differences provide insight into how ECM aging may shape tumor behavior, offering potential targets for modulating the tumor microenvironment across age-related contexts. Given the increasing aged population, strategies that address the ECM directly may enable improvements toward treatment efficacy and metastatic prevention. These findings are consistent with our previous in vitro studies and recent in vivo studies from aged and young mice models.^13,21,22^ By isolating the ECM from senescent cells and mature T and B cells, our model provides evidence that the aged ECM alone can facilitate tumor progression. It enables the study of the ECM alone allowing for the study of both young and aged environment within the same host system, allowing for better opportunities for comparative studies.

## Conclusion

This study demonstrates that the aged ECM alone drives breast cancer growth and progression. Using a novel, hybrid in vivo model, we first find that knockdown of *LOX* reduced tumor volume and suppressed transcriptomic programs linked to metastasis while enriching genes and pathways for ECM structure. *IL1B* and *P4HA1* showed a more complex or potentially context-dependent role in the aged environment, highlighting a need for further studies. We further show that the aged ECM leads to increased tumor growth and reprograms tumor cell states toward invasive and inflammatory phenotypes. Transcriptomic profiling reveals that the aged ECM drives a tumor permissive state while the young promotes a suppressive state. These findings underscore the importance of ECM in cancer progression and therapeutic value and support LOX as a promising therapeutic target. Moreover, this model offers a novel platform for investigating ECM-driven diseases beyond cancer, such as fibrosis and other age-related pathologies.

## Supplementary Figures

**Figure S1:**
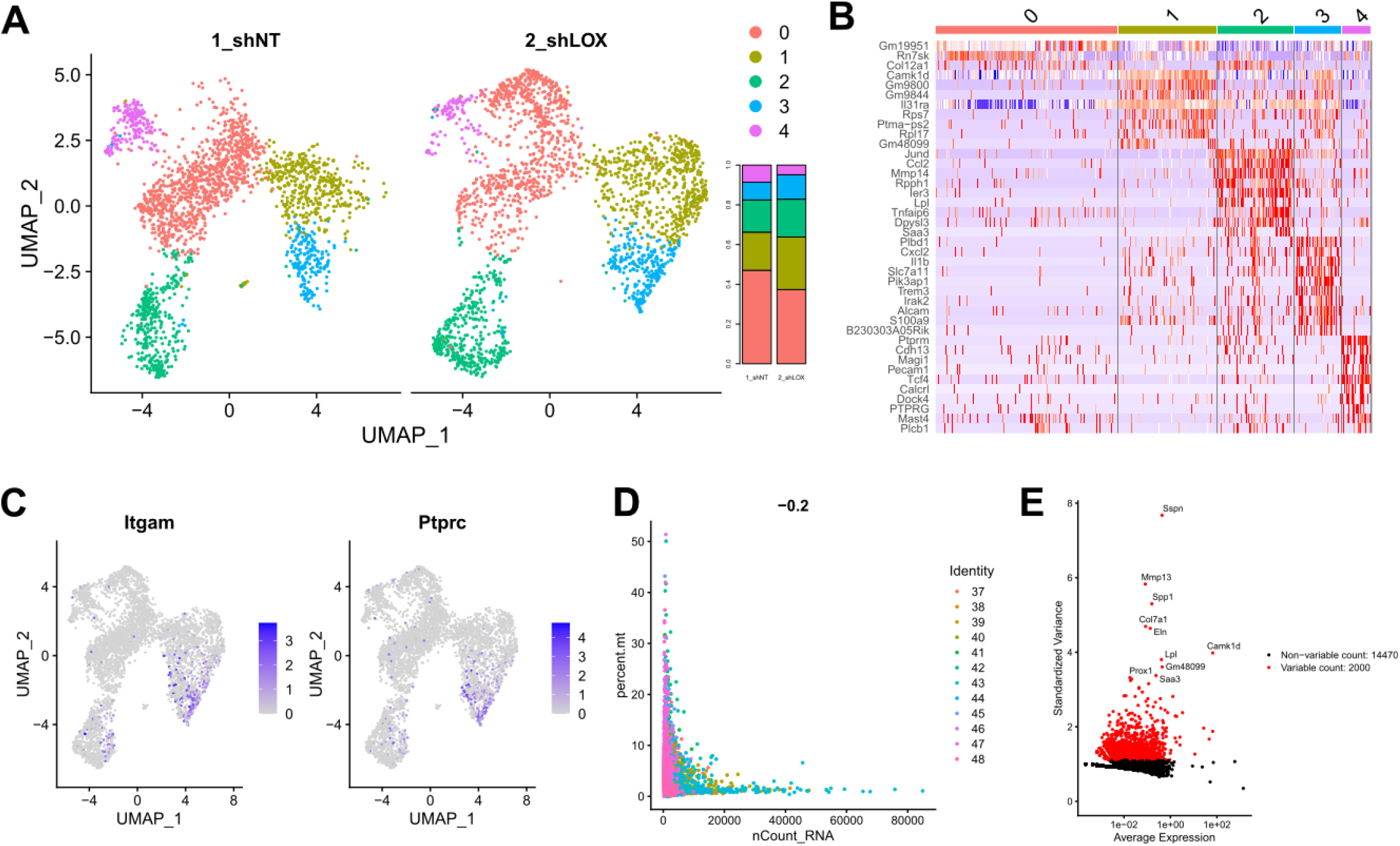
Immune cell group scRNA-seq data. **(A)** Immune cell group UMAP and bar chart depicting the ratio between the clusters in shNT and shLOX groups. **(B)** Immune cell group heat map with marker genes. **(C)** Immune cell group feature plot of Itgam and Ptprc. **(D)** Immune cell group feature scatter plot and **(E)** Variable feature scatter plot.

**Fig S2:**
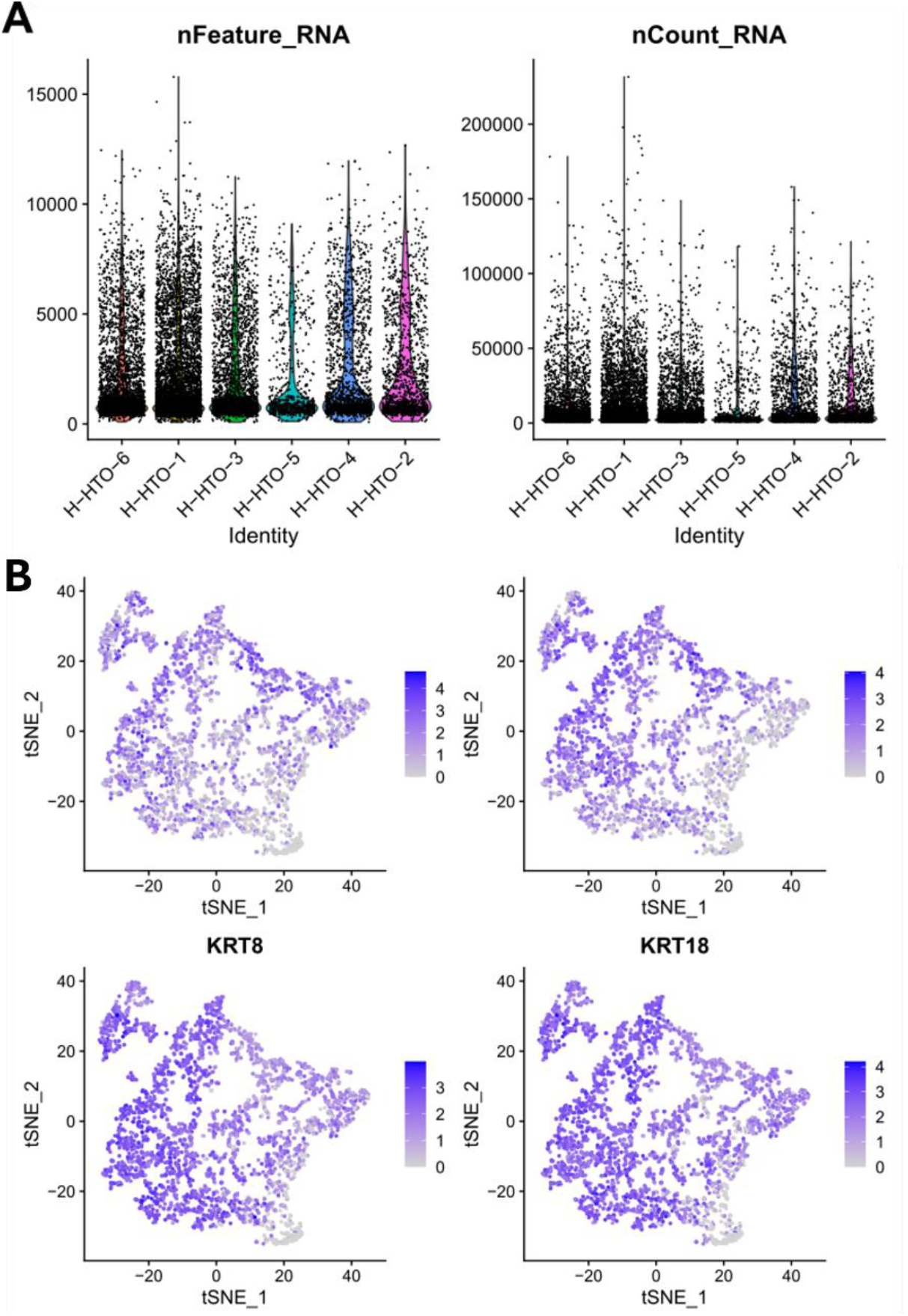
HTO-Breakdown and Keratin Feature Plots of Aged vs. Young ECM Tumors. **(A)** RNA-Feature Chart. **(B)** RNA Count Chart.

**Fig S3:**
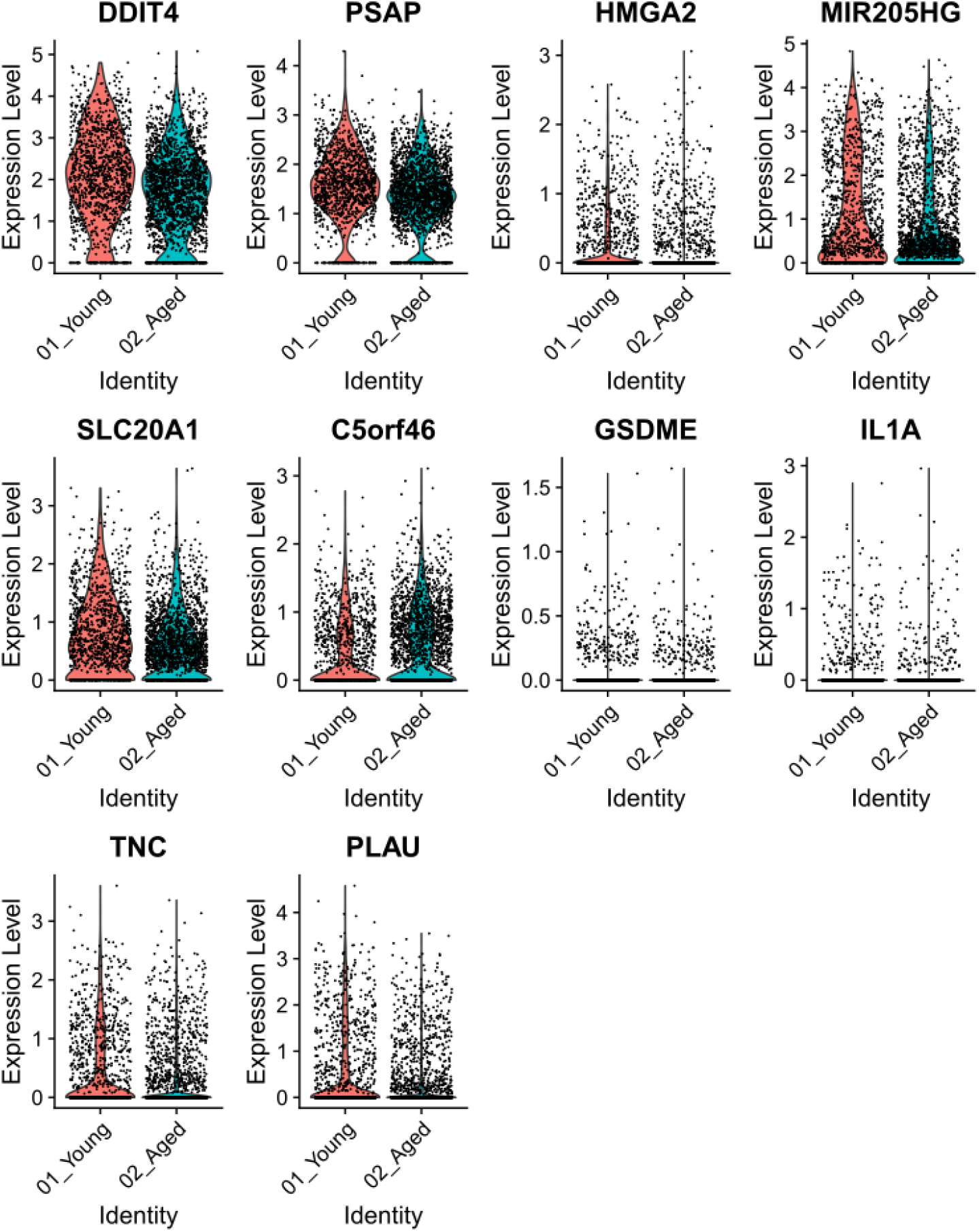
Violin Plots with Significantly Upregulated Genes on the Aged ECM.

## Materials and Methods

### Cell Culturing

GFP tagged MCF10A DCIS.com and all genetic knockdown cells (MCF10A DCIS.com pLKO, MCF10A DCIS.com pLKO-LOX shRNA, MCF10A DCIS.com pLKO-P4HA1 shRNA, and MCF10A DCIS.com pLKO-IL1B shRNA cells) were maintained in DMEM F12 1:1 – DMEM low glucose media in a 1:1 ratio with F12 (Ham’s) media further supplemented with 10% FBS, 1% P/S, and 0.75 µg/mL puromycin.

### Mouse Mammary Breast Removal

4^th^ mammary glands were first isolated from young (2-4 months) and aged (24-48 months) C57BL/6 mice in accordance with IACUC guidelines (protocol number: 18-05-4687) with the approval of Notre Dame, which has an approved Assurance of Compliance on file with the National Institutes of Health, Office of Laboratory Animal Welfare. Mice were sacrificed in CO2 chambers, and tissues collected and used immediately, or wrapped in aluminum foil, flash frozen in liquid nitrogen, and stored at −80 °C until use.

### Decellularization and Delipidation

To section the tissues in cryostat, tissues were thawed at room temperature (RT), blotted on a tissue paper, embedded in optimum cutting temperature (O.C.T.) compound (Tissue-Tek, Sakura, USA), frozen at −20 °C, and sectioned at 300 μm thickness. Sections were washed with PBS to remove the optimum cutting temperature (O.C.T) compound.

For decellularization, whole tissues or tissue sections were incubated in 0.5% SDS for 4 and 2 days, respectively, at 4 °C, with gentle agitation and SDS change every 12 h. For delipidation, isopropanol was used again for 4 and 2 days, respectively, at 4 °C. Tissues and sections (final dimensions ≈2 mm × 2 mm × 0.3 mm) were washed with PBS and stored at 4 °C until use (within 2 weeks).

DNA content of the matrices was measured using the PicoGreen assay as described previously. Briefly, samples (n = 3) were frozen at −80 °C, lyophilized, weighed, incubated in pepsin (Sigma, USA) solution (1 mg mL–1) for 16 h, and the supernatants were assayed in reference to double stranded DNA standards. Results were normalized to dry weights.

### Reseeding of Cells

Green Fluorescent Protein (GFP)-tagged MCF10A DCIS.com (Comedo Ducal Carcinoma In-Situ) cells were then seeded atop the prepared dECM, either young or aged. The reseeded dECM was cultured for 12 days to allow for cellular adhesion, proliferation, and interaction with the ECM. To ensure consistency, 10 similarly sized dECM from each group were selected before implantation. Additionally, the matrices were examined under a fluorescent microscope to validate the similar cellular reseeding. Chosen matrices were completely covered by the GFP-tagged cells.

### Tumor growth and Isolation

Reseeded tissues were implanted into Rag1^-/-^ mice and were inserted into a pocket created between the skin and the peritoneal lining near the 4th mammary fat pad. Weekly, the tumor sizes were recorded via a micrometer, and the mice were sacrificed at 4 weeks (first in vivo) or 6 weeks (second in vivo). The tumors were then excised, dimensions measured using a micrometer, and their volumes calculated.

### Genetic knockdown

To knockdown the IL1B, LOX, and P4HA1 genes, E. coli bacterial stocks containing pLKO.1-Puro plasmids were purchased with the Mission® shRNAs (Sigma-Aldrich) for:

IL1B:

TRCN0000358600 **(IL1B 0)**

TRCN0000358601 **(IL1B 1)**

TRCN0000358661 **(IL1B 61)**

TRCN0000358662 **(IL1B 62)**

LOX:

TRCN0000045989 **(LOX 89)**

TRCN0000045990 **(LOX 90)**

TRCN0000045991 **(LOX 91)**

P4HA1:

TRCN0000303933 **(P4HA1 33)**

TRCN0000303934 **(P4HA1 34)**

TRCN0000303872 **(P4HA1 72)**

Bacteria was cultured in broth culture and plasmids were isolated using the PureLink™ HiPure Plasmid Midiprep Kit (Thermo Fisher Scientific, Cat. No: K210004) following the manufacturer’s instructions.^111^ Briefly, columns were equilibrated using a 10 mL equilibration buffer. In the meanwhile, bacteria (all 10 of them) were spun down in a 50 mL tube at 4000g for 10 min and the pellet was resuspended in 4 mL resuspension buffer, followed by addition of 4 mL of lysis buffer (wait for 5 min) and then 4 mL of precipitation buffer. Tubes were capped and mixed by inverting slowly. Tubes were centrifuged at 13,000g for 10 min. Supernatant was collected and loaded into equilibrated columns, the elute was allowed to drain and the column was washed with Wash buffer (2 times). The elute was discarded. Next, the 5 mL of Elution buffer was added into the column and eluted into sterile 15-mL centrifuge tubes. Isopropanol (3.5 mL) was added, and the tubes were mixed thoroughly. The tube was incubated overnight at 4°C, then centrifuged at 13,000g for 30 min at 4°C, and the pellet was collected and resuspended in 3 mL 70% ethanol. The tubes were centrifuged at 13,000g for 5 min at 4°C and supernatant discarded. Pellet was airdried and resuspended in 200 µL TE buffer. Plasmid DNA amounts were quantified using Nanodrop.

For transfection, plasmids with the specified shRNAs (IL1B 00, IL1B 01, IL1B 61, IL1B62; LOX 89, LOX 90, LOX 91; P4HA1 33, P4HA1 34, P4HA1 72), as well as the control pLKO.1-Puro plasmid with no shRNA, were transfected to HEK293T cells using the Lipofectamine™ 3000 Transfection Reagent (Thermo Scientific, Cat. No: L3000008). Briefly, Lipofectamine 3000 reagent (7.5 µL) was added onto 125 µL Opti-MEM reduced serum medium (Gibco) in a 1 mL centrifuge tube, and plasmid DNA (2.5 µg) containing the specified shRNA and P3000 reagent (5 µL) were added onto 125 µL Opti-MEM medium in a separate 1-mL centrifuge tube.^112^ The solutions were then mixed and incubated for 15 min. Next, the solution (250 µL) was added onto HEK293T cells plated in the well of a 6-well plate (at around 70% confluency) containing 2 mL growth medium (DMEM with 10% FBS). Medium was refreshed after 24 h to remove the plasmid DNA-Lipofectamine in the medium. The medium containing the lentiviral particles (1.5 to 2 mL) was collected at days 2 and 4 after transfection. The medium was added onto MCF10A DCIS.com cells plated in 75-mL culture flasks and containing 10 mL growth medium (DMEM-F12 (1:1) with 10% FBS). This was done with both day 2 and day 4 supernatants collected from HEK cells separately (2 times) after refreshing the DMEM-F12 media on DCIS.com cells. After a week, the DCIS.com cell were treated with growth medium containing 0.75 µg/mL puromycin to select the transduced cells. After selection, cells were replated and used in future experiments.

### Western Blot

Knockdown DCIS.com cells were replated in 12-well plates and cultured for 5 days. Next, the cells were washed and lysed using 200 µL RIPA buffer (30 min in 4°C). Protein amounts were quantified using BCA assay, and lysates containing 15 µg of proteins were loaded into SDS-PAGE. Before loading the samples, loading buffer (4x) and β-mercaptoethanol were added onto the samples. Samples were run in electrophoresis and transferred to nitrocellulose membranes as described previously.^13^ For staining, the following antibodies were used:

Beta Actin (Abcam, rb pAb to Beta Actin, Cat No: ab8227), 1:1000 dilution

IL1B antibody (Cell Signaling, IL-1b (D3U3E) Rabbit mAb, Cat. No: 12703S), 1:1000 dilution) LOX antibody (Abcam, Rabbit Recombinant Anti-LOX antibody [EPR4025], Cat. No: ab174316), 1:400 dilution)

P4HA1 antibody (Abcam, Anti-P4HA1 rabbit antibody, Cat. No: ab244400), 1:1000 dilution) Secondary Antibody (Abcam, Goat Anti-Rabbit IgG H&L (HRP), Cat No: ab6721), 1:2000 dilution)

The MCF10A DCIS.com cells transduced with IL1B 62, LOX 91, and P4HA1 33 shRNAs were selected, as they yielded the best knockdown efficiency.

### scRNA-seq of Models with Control and Knockdown Cells

Instead of the cells being labelled with human HTO, CD11b was used to separate the immune cells (CD11b+) from the tumor cells (CD11b-) (Milenyi Biotec, CD11b MicroBeads UltraPure, Mouse Cat. No: 130-126-725). As the tumor cells were human in origin, the bead separation ensured the enrichment of the tumor cells so that it can be fully aligned to the readout of the human genome. Cells were fixed, barcoded, and subjected to library preparation for next-generation sequencing. Cells were then fixed using the Evercode Cell Fixation v2 kit (Cat. No: ECF2101) following the Parse Fixation Protocol^113^. Fixed cells were shipped on dry ice to UT Southwestern Medical Center for library preparation and preserved at –80 ºC. Next, the fixed cells underwent three rounds of barcoding using the Parse Bioscience Evercode whole transcriptome WT v2 kit with a target recovery of 50,000 cells total, followed by cell lysis, and sublibrary preparation. 8 sublibraries were sequenced on a NovaSeq-X instrument and the raw data was processed using the Parse Biosciences analysis pipeline. Reads obtained from the tumor cell samples were aligned to the human reference genome while the immune (CD11b+) sample reads were aligned to the mouse genome. Downstream processing, RNA normalization, and DEG analysis was performed in RStudio using the Seurat package.

### scRNA-seq of Models with Young and Aged dECM

Three sets of tumors (6 young and 6 aged) were selected for single-cell RNA-sequencing (scRNA-seq). To process the tumors for scRNAseq, tumors were minced using blades, and the matrix was disassociated using 5 mL enzyme mix (Milenyi Biotec, Human Tumor Dissociation Kit Cat. No: 130-095-929) for each tumor in a 50 mL tube. Mixtures were incubated at 37°C with continuous rotation for 50 min. Undissolved tissue was removed using cell strainers (pores size: 70 µm) and DMEM media was passed (20 mL) through the strainers. The cell suspension was centrifuged at 300g for 7 min and supernatant was aspirated. The pellet was reconstituted in DMEM media and cell aggregates were removed by straining the cells through 30 µm strainers. Next, dead cell removal kit (Miltenyi, Cat. No: 130-090-101) was used to remove the dead cells. The cells were diluted with DMEM media and tubes were centrifuged. Cells were reconstituted in dead cell removal microbeads (100 µL per 10^7^ cells) and incubated for 15 min at room temperature. Next, 500 µL of buffer was added on to the cells and the cell suspension was poured through equilibrated columns by placing on the magnets on a MACS separator. Effluent (live cell fraction) was collected, and the cells were labelled with human-HTO antibodies and sent to 10x Genomics for analysis.

## References

1. Sung, H. et al. Global Cancer Statistics 2020: GLOBOCAN Estimates of Incidence and Mortality Worldwide for 36 Cancers in 185 Countries. CA: A Cancer Journal for Clinicians 71, 209–249 (2021).

2. Arnold, M. et al. Current and future burden of breast cancer: Global statistics for 2020 and 2040. The Breast 66, 15–23 (2022).

3. Lei, S. et al. Global patterns of breast cancer incidence and mortality: A population-based cancer registry data analysis from 2000 to 2020. Cancer Communications 41, 1183–1194 (2021).

4. Key, T. J., Verkasalo, P. K. & Banks, E. Epidemiology of breast cancer. The Lancet Oncology 2, 133–140 (2001).

5. Bray, F. et al. Global cancer statistics 2022: GLOBOCAN estimates of incidence and mortality worldwide for 36 cancers in 185 countries. CA: A Cancer Journal for Clinicians 74, 229–263 (2024).

6. Sun, Y.-S. et al. Risk Factors and Preventions of Breast Cancer. Int J Biol Sci 13, 1387–1397 (2017).

7. López-Otín, C., Pietrocola, F., Roiz-Valle, D., Galluzzi, L. & Kroemer, G. Meta-hallmarks of aging and cancer. Cell Metabolism 35, 12–35 (2023).

8. Klement, R. J. Cancer as a global health crisis with deep evolutionary roots. Global Transitions 6, 45–65 (2024).

9. Tamayo-Angorrilla, M., López de Andrés, J., Jiménez, G. & Marchal, J. A. The biomimetic extracellular matrix: a therapeutic tool for breast cancer research. Translational Research 247, 117–136 (2022).

10. Owyong, M. et al. Overcoming Barriers of Age to Enhance Efficacy of Cancer Immunotherapy: The Clout of the Extracellular Matrix. Frontiers in Cell and Developmental Biology 6, (2018).

11. Gosselin, K. et al. Senescence-Associated Oxidative DNA Damage Promotes the Generation of Neoplastic Cells. Cancer Research 69, 7917–7925 (2009).

12. Sprenger, C. C., Plymate, S. R. & Reed, M. J. Aging-related alterations in the extracellular matrix modulate the microenvironment and influence tumor progression. International Journal of Cancer 127, 2739–2748 (2010).

13. Bahcecioglu, G. et al. Aged Breast Extracellular Matrix Drives Mammary Epithelial Cells to an Invasive and Cancer-Like Phenotype. Adv Sci (Weinh) 8, 2100128 (2021).

14. Yang, J., Bahcecioglu, G., Ronan, G. & Zorlutuna, P. Aged breast matrix bound vesicles promote breast cancer invasiveness. Biomaterials 306, 122493 (2024).

15. Yousefzadeh, M. J. et al. An aged immune system drives senescence and ageing of solid organs. Nature 594, 100–105 (2021).

16. Weyand, C. M. & Goronzy, J. J. Aging of the Immune System. Mechanisms and Therapeutic Targets. Ann Am Thorac Soc 13, S422–S428 (2016).

17. Full article: N-acetyl cysteine ameliorates aortic fibrosis by promoting M2 macrophage polarization in aging mice. https://www.tandfonline.com/doi/full/10.1080/13510002.2021.1976568#abstract.

18. Pan, Y., Yu, Y., Wang, X. & Zhang, T. Tumor-Associated Macrophages in Tumor Immunity. Front Immunol 11, 583084 (2020).

19. Zhang, C., Gu, X., Pan, M., Yuan, Q. & Cheng, H. Senescent thyroid tumor cells promote their migration by inducing the polarization of M2-like macrophages. Clin Transl Oncol 23, 1253–1261 (2021).

20. Li, Y. et al. Age-related macrophage alterations are associated with carcinogenesis of colorectal cancer. Carcinogenesis 43, 1039–1049 (2022).

21. Angarola, B. L. et al. Comprehensive single-cell aging atlas of healthy mammary tissues reveals shared epigenomic and transcriptomic signatures of aging and cancer. Nat Aging 5, 122–143 (2025).

22. Falvo, P. et al. Age-dependent differences in breast tumor microenvironment: challenges and opportunities for efficacy studies in preclinical models. Cell Death Differ 1–14 (2025) doi:10.1038/s41418-025-01447-1.

23. Statzer, C., Park, J. Y. C. & Ewald, C. Y. Extracellular Matrix Dynamics as an Emerging yet Understudied Hallmark of Aging and Longevity. Aging Dis 14, 670–693 (2023).

24. Strategic outline of interventions targeting extracellular matrix for promoting healthy longevity | American Journal of Physiology-Cell Physiology. https://journals.physiology.org/doi/full/10.1152/ajpcell.00060.2023.

25. Jackson, S. J. et al. Does age matter? The impact of rodent age on study outcomes. Lab Anim 51, 160–169 (2017).

26. Flynn, J. N., Horan, L., Tucker, C. S., Robb, D. & Wilkinson, M. J. A. 3Rs considerations when using ageing animals in science. in The UFAW Handbook on the Care and Management of Laboratory and Other Research Animals 251–267 (John Wiley & Sons, Ltd, 2024). doi:10.1002/9781119555278.ch16.

27. Sun, M. et al. The need to incorporate aged animals into the preclinical modeling of neurological conditions. Neuroscience & Biobehavioral Reviews 109, 114–128 (2020).

28. Lian, J., Yue, Y., Yu, W. & Zhang, Y. Immunosenescence: a key player in cancer development. J Hematol Oncol 13, 151 (2020).

29. Ecker, B. L. et al. Age-Related Changes in HAPLN1 Increase Lymphatic Permeability and Affect Routes of Melanoma Metastasis. Cancer Discov 9, 82–95 (2019).

30. Sceneay, J. et al. Interferon Signaling Is Diminished with Age and Is Associated with Immune Checkpoint Blockade Efficacy in Triple-Negative Breast Cancer. Cancer Discovery 9, 1208–1227 (2019).

31. Toney, N. J. et al. B cells enhance IL-1 beta driven invasiveness in triple negative breast cancer. Sci Rep 15, 2211 (2025).

32. Vikhreva, P. et al. TAp73 upregulates IL-1β in cancer cells: Potential biomarker in lung and breast cancer? Biochemical and Biophysical Research Communications 482, 498–505 (2017).

33. Xu, R. P4HA1 is a new regulator of the HIF-1 pathway in breast cancer. Cell Stress 3, 27–28.

34. Xiong, G. et al. Collagen prolyl 4-hydroxylase 1 is essential for HIF-1α stabilization and TNBC chemoresistance. Nat Commun 9, 4456 (2018).

35. Li, J. et al. Lysyl Oxidase (LOX) Family Proteins: Key Players in Breast Cancer Occurrence and Progression. J Cancer 15, 5230–5243 (2024).

36. Homo sapiens (human) long intergenic non-protein coding RNA 1091 (LINC01091) | URS0000CCE120. https://rnacentral.org/rna/URS0000CCE120/9606.

37. LINC01091 long intergenic non-protein coding RNA 1091 [Homo sapiens (human)] - Gene - NCBI. https://www.ncbi.nlm.nih.gov/gene?Db=gene&Cmd=DetailsSearch&Term=285419.

38. Liu, K., Zhang, L., Li, X. & Zhao, J. High expression of lncRNA <em>HSD11B1-AS1</em> indicates favorable prognosis and is associated with immune infiltration in cutaneous melanoma. Oncology Letters 23, 1–14 (2022).

39. HSD11B1-AS1 HSD11B1 antisense RNA 1 [Homo sapiens (human)] - Gene - NCBI. https://www.ncbi.nlm.nih.gov/gene/101930114.

40. Xu, Z. et al. Comprehensive Analysis of Ferroptosis-Related LncRNAs in Breast Cancer Patients Reveals Prognostic Value and Relationship With Tumor Immune Microenvironment. Front. Surg. 8, (2021).

41. ABCC2 gene: MedlinePlus Genetics. https://medlineplus.gov/genetics/gene/abcc2/.

42. TMCC3 transmembrane and coiled-coil domain family 3 [Homo sapiens (human)] - Gene - NCBI. https://www.ncbi.nlm.nih.gov/gene/57458.

43. Wang, Y.-H. et al. Transmembrane and coiled-coil domain family 3 (TMCC3) regulates breast cancer stem cell and AKT activation. Oncogene 40, 2858–2871 (2021).

44. MUC5B mucin 5B, oligomeric mucus/gel-forming [Homo sapiens (human)] - Gene - NCBI. https://www.ncbi.nlm.nih.gov/gene/727897.

45. MUC16 mucin 16, cell surface associated [Homo sapiens (human)] - Gene - NCBI. https://www.ncbi.nlm.nih.gov/gene/94025.

46. Zhang, X.-Y.Hong, L.-L. & Ling, Z. MUC16: clinical targets with great potential. Clin Exp Med 24, 101 (2024).

47. MMP7 matrix metallopeptidase 7 [Homo sapiens (human)] - Gene - NCBI. https://www.ncbi.nlm.nih.gov/gene/4316.

48. RASA3 RAS p21 protein activator 3 [Homo sapiens (human)] - Gene - NCBI. https://www.ncbi.nlm.nih.gov/gene/22821.

49. Johansen, K. H., Golec, D. P., Okkenhaug, K. & Schwartzberg, P. L. Mind the GAP: RASA2 and RASA3 GTPase-activating proteins as gatekeepers of T cell activation and adhesion. Trends in Immunology 44, 917–931 (2023).

50. MMP3 matrix metallopeptidase 3 [Homo sapiens (human)] - Gene - NCBI. https://www.ncbi.nlm.nih.gov/gene/4314.

51. HLA-F-AS1 HLA-F antisense RNA 1 [Homo sapiens (human)] - Gene - NCBI. https://www.ncbi.nlm.nih.gov/gene?Db=gene&Cmd=DetailsSearch&Term=285830.

52. Huang, Y. et al. HLA-F-AS1/miR-330-3p/PFN1 axis promotes colorectal cancer progression. Life Sciences 254, 117180 (2020).

53. Zhang, X., Li, W., Li, S., Zhang, Z. & Song, W. Potentials as biomarker and therapeutic target of upregulated long non-coding RNA HLA-F antisense RNA 1 in hepatitis B virus-associated hepatocellular carcinoma. Virus Genes 60, 243–250 (2024).

54. HIF1A-AS3 HIF1A antisense RNA 3 [Homo sapiens (human)] - Gene - NCBI. https://www.ncbi.nlm.nih.gov/gene/105370526.

55. Xie, W. et al. A novel hypoxia-stimulated lncRNA HIF1A-AS3 binds with YBX1 to promote ovarian cancer tumorigenesis by suppressing p21 and AJAP1 transcription. Molecular Carcinogenesis 62, 1860–1876 (2023).

56. Mansoori, B. et al. HMGA2 as a Critical Regulator in Cancer Development. Genes (Basel) 12, 269 (2021).

57. Calcium-Activated Potassium Channel (KCNMA1) as Biomarker of Pre-Invasive and Invasive Cervical Cancer | Indian Journal of Gynecologic Oncology. https://link.springer.com/article/10.1007/s40944-024-00822-z.

58. The JAK/STAT signaling pathway: from bench to clinic | Signal Transduction and Targeted Therapy. https://www.nature.com/articles/s41392-021-00791-1.

59. Choi, R. B. & Robling, A. G. The Wnt pathway: an important control mechanism in bone’s response to mechanical loading. Bone 153, 116087 (2021).

60. Piccolo, S., Panciera, T., Contessotto, P. & Cordenonsi, M. YAP/TAZ as master regulators in cancer – modulation, function and therapeutic approaches. Nat Cancer 4, 9–26 (2023).

61. Mehrgou, A. & Akouchekian, M. The importance of BRCA1 and BRCA2 genes mutations in breast cancer development. Med J Islam Repub Iran 30, 369 (2016).

62. Karakostis, K., Malbert-Colas, L., Thermou, A., Vojtesek, B. & Fåhraeus, R. The DNA damage sensor ATM kinase interacts with the p53 mRNA and guides the DNA damage response pathway. Molecular Cancer 23, 21 (2024).

63. KRT23 keratin 23 [Homo sapiens (human)] - Gene - NCBI. https://www.ncbi.nlm.nih.gov/gene/25984.

64. Zhang, N. et al. Keratin 23 promotes telomerase reverse transcriptase expression and human colorectal cancer growth. Cell Death Dis 8, e2961–e2961 (2017).

65. DNER delta/notch like EGF repeat containing [Homo sapiens (human)] - Gene - NCBI. https://www.ncbi.nlm.nih.gov/gene/92737.

66. DNER promotes epithelial–mesenchymal transition and prevents chemosensitivity through the Wnt/β-catenin pathway in breast cancer | Cell Death & Disease. https://www.nature.com/articles/s41419-020-02903-1.

67. TSPAN2 tetraspanin 2 [Homo sapiens (human)] - Gene - NCBI. https://www.ncbi.nlm.nih.gov/gene/10100.

68. Zhang, H. et al. Tspan protein family: focusing on the occurrence, progression, and treatment of cancer. Cell Death Discov. 10, 187 (2024).

69. CEP55 centrosomal protein 55 [Homo sapiens (human)] - Gene - NCBI. https://www.ncbi.nlm.nih.gov/gene/55165.

70. Jeffery, J., Sinha, D., Srihari, S., Kalimutho, M. & Khanna, K. K. Beyond cytokinesis: the emerging roles of CEP55 in tumorigenesis. Oncogene 35, 683–690 (2016).

71. HMMR hyaluronan mediated motility receptor [Homo sapiens (human)] - Gene - NCBI. https://www.ncbi.nlm.nih.gov/gene/3161.

72. Mateo, F. et al. Modification of BRCA1-associated breast cancer risk by HMMR overexpression. Nat Commun 13, 1895 (2022).

73. CCNA2 cyclin A2 [Homo sapiens (human)] - Gene - NCBI. https://www.ncbi.nlm.nih.gov/gene/890.

74. Gan, Y., Li, Y., Li, T., Shu, G. & Yin, G. CCNA2 acts as a novel biomarker in regulating the growth and apoptosis of colorectal cancer. Cancer Manag Res 10, 5113–5124 (2018).

75. KIF2C kinesin family member 2C [Homo sapiens (human)] - Gene - NCBI. https://www.ncbi.nlm.nih.gov/gene/11004.

76. Li, R.-Q. et al. KIF2C: An important factor involved in signaling pathways, immune infiltration, and DNA damage repair in tumorigenesis. Biomedicine & Pharmacotherapy 171, 116173 (2024).

77. KIF18B kinesin family member 18B [Homo sapiens (human)] - Gene - NCBI. https://www.ncbi.nlm.nih.gov/gene/146909.

78. Wu, Y.-P. et al. Kinesin family member 18B regulates the proliferation and invasion of human prostate cancer cells. Cell Death Dis 12, 302 (2021).

79. MCM6 minichromosome maintenance complex component 6 [Homo sapiens (human)] - Gene - NCBI. https://www.ncbi.nlm.nih.gov/gene/4175.

80. Zeng, T. et al. The DNA replication regulator MCM6: An emerging cancer biomarker and target. Clinica Chimica Acta 517, 92–98 (2021).

81. E2F1 E2F transcription factor 1 [Homo sapiens (human)] - Gene - NCBI. https://www.ncbi.nlm.nih.gov/gene/1869.

82. Ashok, C. et al. E2F1 and epigenetic modifiers orchestrate breast cancer progression by regulating oxygen-dependent ESRP1 expression. Oncogenesis 10, 58 (2021).

83. WDR76 WD repeat domain 76 [Homo sapiens (human)] - Gene - NCBI. https://www.ncbi.nlm.nih.gov/gene/79968.

84. Jeong, W.-J. et al. WDR76 is a RAS binding protein that functions as a tumor suppressor via RAS degradation. Nat Commun 10, 295 (2019).

85. DTL denticleless E3 ubiquitin protein ligase adapter [Homo sapiens (human)] - Gene - NCBI. https://www.ncbi.nlm.nih.gov/gene/51514.

86. Liu, S. et al. Overexpression of DTL enhances cell motility and promotes tumor metastasis in cervical adenocarcinoma by inducing RAC1-JNK-FOXO1 axis. Cell Death Dis 12, 929 (2021).

87. KCNMA1 potassium calcium-activated channel subfamily M alpha 1 [Homo sapiens (human)] - Gene - NCBI. https://www.ncbi.nlm.nih.gov/gene/3778.

88. Ma, G. et al. KCNMA1 cooperating with PTK2 is a novel tumor suppressor in gastric cancer and is associated with disease outcome. Molecular Cancer 16, 46 (2017).

89. TRPV3 - Gene - NCBI. https://www.ncbi.nlm.nih.gov/gene/?term=TRPV3.

90. Xie, Y., Kim, H. I., Yang, Q., Wang, J. & Huang, W. TRPV3 regulates Breast Cancer Cell Proliferation and Apoptosis by EGFR/AKT pathway. J Cancer 15, 2891–2899 (2024).

91. PLCG2 phospholipase C gamma 2 [Homo sapiens (human)] - Gene - NCBI. https://www.ncbi.nlm.nih.gov/gene/5336.

92. Zhao, Y. et al. Overexpression of PLCG2 and TMEM38A inhibit tumor progression in clear cell renal cell carcinoma. Sci Rep 15, 3192 (2025).

93. C6orf141 chromosome 6 open reading frame 141 [Homo sapiens (human)] - Gene - NCBI. https://www.ncbi.nlm.nih.gov/gene/135398.

94. Yang, C.-M. et al. Low C6orf141 Expression is Significantly Associated with a Poor Prognosis in Patients with Oral Cancer. Sci Rep 9, 4520 (2019).

95. MMP1 matrix metallopeptidase 1 [Homo sapiens (human)] - Gene - NCBI. https://www.ncbi.nlm.nih.gov/gene/4312.

96. MMP9 matrix metallopeptidase 9 [Homo sapiens (human)] - Gene - NCBI. https://www.ncbi.nlm.nih.gov/gene/4318.

97. MMP10 matrix metallopeptidase 10 [Homo sapiens (human)] - Gene - NCBI. https://www.ncbi.nlm.nih.gov/gene/4319.

98. Dai, L. et al. Comprehensive bioinformatic analysis of MMP1 in hepatocellular carcinoma and establishment of relevant prognostic model. Sci Rep 12, 13639 (2022).

99. Li, F., Chen, L., Xia, Q., Feng, Z. & Li, N. Combined knockdown of CD151 and MMP9 may inhibit the malignant biological behaviours of triple-negative breast cancer through the GSK-3β/β-catenin-related pathway. Sci Rep 14, 21786 (2024).

100. Zhang, G., Miyake, M., Lawton, A., Goodison, S. & Rosser, C. J. Matrix metalloproteinase-10 promotes tumor progression through regulation of angiogenic and apoptotic pathways in cervical tumors. BMC Cancer 14, 310 (2014).

101. Dharavath, B. et al. Role of miR-944/MMP10/AXL- axis in lymph node metastasis in tongue cancer. Commun Biol 6, 57 (2023).

102. CLCA2 chloride channel accessory 2 [Homo sapiens (human)] - Gene - NCBI. https://www.ncbi.nlm.nih.gov/gene/9635.

103. Zhang, P., Lin, Y. & Liu, Y. CLCA2 suppresses the proliferation, migration and invasion of cervical cancer. Exp Ther Med 22, 776 (2021).

104. CADM3 cell adhesion molecule 3 [Homo sapiens (human)] - Gene - NCBI. https://www.ncbi.nlm.nih.gov/gene/57863.

105. Ren, H. et al. Clinical significance of low expression of CADM3 in breast cancer and preliminary exploration of related mechanisms. BMC Cancer 24, 367 (2024).

106. DDIT4 DNA damage inducible transcript 4 [Homo sapiens (human)] - Gene - NCBI. https://www.ncbi.nlm.nih.gov/gene/54541.

107. Chen, X. et al. Identification of DDIT4 as a potential prognostic marker associated with chemotherapeutic and immunotherapeutic response in triple-negative breast cancer. World J Surg Oncol 21, 194 (2023).

108. Tajik, F. et al. Nuclear overexpression of DNA damage-inducible transcript 4 (DDIT4) is associated with aggressive tumor behavior in patients with pancreatic tumors. Sci Rep 13, 19403 (2023).

109. C5orf46 chromosome 5 open reading frame 46 [Homo sapiens (human)] - Gene - NCBI. https://www.ncbi.nlm.nih.gov/gene/389336.

110. Ma, M. et al. Preliminary study on the role of the C5orf46 gene in renal cancer. Translational Oncology 21, 101442 (2022).

111. TFS-AssetsLSGmanualspurelink_hipure_plasmid_dna_purification_man.pdf.

112. Protocol for cell preparation and gene delivery in HEK293T and C2C12 cells - ScienceDirect. https://www.sciencedirect.com/science/article/pii/S2666166721002045?via%3Dihub.

113. Evercode Cell Fixation v3 User Guides. Support Suite - Parse Biosciences https://support.parsebiosciences.com/hc/en-us/articles/23914547728660-Evercode-Cell-Fixation-v3-User-Guides (2024).

